# Generation and evaluation of protease inhibitor-resistant SARS-CoV-2 strains

**DOI:** 10.1101/2023.11.22.568013

**Authors:** Hawa Sophia Bouzidi, Jean-Sélim Driouich, Raphaëlle Klitting, Ornéllie Bernadin, Géraldine Piorkowski, Rayane Amaral, Laurent Fraisse, Charles E. Mowbray, Ivan Scandale, Fanny Escudié, Eric Chatelain, Xavier de Lamballerie, Antoine Nougairède, Franck Touret

## Abstract

Since the start of the SARS-CoV-2 pandemic, the search for antiviral therapies has been at the forefront of medical research. To date, the 3CLpro inhibitor nirmatrelvir (Paxlovid®) has shown the best results in clinical trials and the greatest robustness against variants. A second SARS-CoV-2 protease inhibitor, ensitrelvir (Xocova®), has been developed. Ensitrelvir, currently in Phase 3, was approved in Japan under the emergency regulatory approval procedure in November 2022, and is available since March 31, 2023. One of the limitations for the use of antiviral monotherapies is the emergence of resistance mutations. Here, we experimentally generated mutants resistant to nirmatrelvir and ensitrelvir *in vitro* following repeating passages of SARS-CoV-2 in the presence of both antivirals. For both molecules, we demonstrated a loss of sensitivity for resistance mutants *in vitro*. Using a Syrian golden hamster infection model, we showed that the ensitrelvir M49L mutation, in the multi-passage strain, confers a high level of *in vivo* resistance. Finally, we identified a recent increase in the prevalence of M49L-carrying sequences, which appears to be associated with multiple repeated emergence events in Japan and may be related to the use of Xocova® in the country since November 2022. These results highlight the strategic importance of genetic monitoring of circulating SARS-CoV-2 strains to ensure that treatments administered retain their full effectiveness.

## Introduction

Since the beginning of the SARS-CoV-2 pandemic in 2020, the search for antiviral therapeutics has been a major focus for medical research. As previously observed with Influenza and HIV, effective countermeasures to treat people can be decisive in managing a health crisis. At the start of the pandemic, the repositioning of broad-spectrum antivirals targeting the replication complex raised great expectations(Wang et al., 2020; Williamson et al., 2020), which were dashed by disappointing results in clinical trials(Beigel et al., 2020). Therapeutic antibodies have subsequently demonstrated their usefulness(Pinto et al., 2020; Starr et al., 2021), but failed to overcome the emergence of variants escaping the humoral response(Touret et al., 2023). Ultimately, the best effects in clinical trials(Hammond et al., 2022) and the greatest robustness against variants(Vangeel et al., 2022) were achieved with the protease inhibitor nirmatrelvir(Owen et al., 2021) (PF-07321332, Paxlovid®), targeting the 3CLpro. This inhibitor of the coronavirus protease 3CLpro (PF-00835231) was originally developed against SARS-CoV-1 (Boras et al., 2021). It is also effective against SARS-CoV-2 by preventing the cleavage of the polyproteins PP1a and PP1ab – effectively blocking the generation of the non-structural proteins essential for viral replication(Jin et al., 2020).

The pharmacokinetics (PK) of nirmatrelvir, whose metabolism includes a predominant role for CYP3A4(Owen et al., 2021), have been improved by the combination with ritonavir used as a exposure booster. However, this combination may complicate the use of nimatrelvir in certain patients receiving other drugs whose metabolism may also be modified by ritonavir(Hoertel et al., 2022).

In the meantime, a second SARS-CoV-2 3CLpro inhibitor named ensitrelvir (S-217622), was developed by the Japanese company Shionogi. The molecule was identified through a combination of virtual and biological screenings followed by optimization using a structure-based drug design strategy(Unoh et al., 2022). It has been proposed for use without an association with ritonavir(Unoh et al., 2022). Following encouraging results in Phase 2 clinical trials, it is now in Phase 3(Shionogi, 2023; University of Minnesota, 2023). In Japan, however, ensitrelvir (Xocova®), was approved under the emergency regulatory approval procedure in November 2022 and has been available on prescription since March 31, 2023.

Resistance mutations can represent a major limitation to the use of antiviral mono-therapies. Such mutations may naturally be observed in a proportion of circulating strains in the absence of any apparent specific selective pressure(Bloom et al., 2010; Ip et al., 2023; Kawashima et al., 2023) and/or appear in patients in the context of conditions of use associated with sub-optimal efficiency (e.g., during prolonged use in a patient with long lasting viral replication). For instance, the emergence of resistance against antiviral monotherapy has been observed in patients treated with the monoclonal antibody sotrovimab during the Delta wave of SARS-CoV-2(Rockett et al., 2022).

Here, we experimentally generated nirmatrelvir and ensitrelvir resistance mutants. For both molecules, we demonstrated a loss in sensitivity for the resistance mutants *in vitro*. Using a Syrian golden hamster model of SARS-CoV-2 infection, we showed that the ensitrelvir resistance mutation M49L confers a high level of resistance *in vivo*. Finally, we identified a recent increase in the prevalence of M49L-carrying sequences, that we link to multiple repeated emergence events in Japan, and may be associated with the commercialization of the molecule in the country since April 2023.

## Materiel and methods

### Experimental model

#### Cell line

VeroE6/TMPRSS2 cells (ID 100978) were obtained from CFAR and were grown in MEM (Minimal Essential Medium-Life Technologies) with 7 .5% heat-inactivated Fetal Calf Serum (FCS; Life Technologies with 1% penicillin/streptomycin PS, 5000U.mL^−1^ and 5000µg.mL^−1^ respectively (Life Technologies) and supplemented with 1 % non-essential amino acids (Life Technologies) and G-418 (Life Technologies), at 37°C with 5% CO_2_.

#### Viral strain

SARS-CoV-2 strain BavPat1 was obtained from Pr. C. Drosten through EVA GLOBAL (https://www.european-virus-archive.com/) and contains the D614G mutation.

#### Organisms/strains

Three-week-old female Syrian hamsters (Mesocricetus auratus) were purchased from Janvier labs. Three-week-old Syrian hamsters (Janvier labs) were maintained in ISOcage P - Bioexclusion System (Techniplast) with unlimited access to water/food and 14h/10h light/dark cycle. Animals were monitored and weighed daily throughout the duration of the study to detect the appearance of any clinical signs of illness/suffering.

##### Antiviral Compounds

Remdesivir was purchased from BLDpharm. ensitrelvir and nirmatrelvir were purchased from MedChemexpress

## Method details

### RNA extraction and quantification

Viral RNA was extracted from 100 μL of cell supernatant from passages P8 and P16 using a QIAamp Viral RNA kit on the automated QIAcube (Qiagen), following the manufacturer’s instructions. Relative quantification of viral RNA was performed using the GoTaq® 1-Step RT-qPCR System kit (Promega). The mixture contained 5 μL of 2x Master Mix, 0.25 μL of each primer (250 nM), 0.07 μL of probe (75 nM), 0.2 μL of GoScript RT Mix and 3.8 μL of extracted nucleic acids. Assays were performed using the QuantStudio 12 K Flex real-time PCR machine (Life technologies) under the following conditions : 50°C for 15 min, 95°C for 2 min, followed by 40 cycles of 95°C for 3 s, 60°C for 30 s. Data collection took place during the 60°C step. Synthetic RNA was used to calculate the amount of viral RNA from standard curves.

### EC_50_ determination

One day prior to infection, 5×10^4^ VeroE6/TMPRSS2 cells per well were seeded in 100µL assay medium (containing 2.5% FBS) in 96 well culture plates.The next day, antiviral compounds were added using the D300e dispenser (TECAN) with eight ½ dilutions. Then, 50 µL/well of a virus mix diluted in medium was added to the wells. Each well was inoculated with 100 TCID_50_ of virus which correspond here to a MOI at 0.002 as classically used for SARS-CoV-2(Touret et al., 2020a). Prior to the assay it was verified for each variant that with this MOI, viruses in the cell culture supernatants were harvested during the logarithmic growth phase of viral replication at 48 hours post infection (Touret et al., 2022, 2019). Four virus control wells were included within the plate. Quantification of the viral genome by real-time RT-qPCR as previously described (Touret et al., 2020b).Nucleic acid from 100µL of cell supernatant were extracted using QIAamp 96 DNA kit and Qiacube HT robot (both from Qiagen). Viral RNA was quantified by real-time RT-qPCR (GoTaq 1 step RT-qPCR kit, Promega). Quantification was provided by serial dilutions of an appropriate T7-generated synthetic RNA standard. RT-qPCR reactions were performed on QuantStudio 12K Flex Real-Time PCR System (Applied Biosystems) and analyzed using QuantStudio 12K Flex Applied Biosystems software v1.2.3. Primers and probe sequences, which target SARS-CoV-2 N gene, were: Fw: 5’-GGCCGCAAATTGCACAAT-3’ ; Rev : 5’-CCAATGCGCGACATTCC-3’; Probe: 5’-FAM-CCCCCAGCGCTTCAGCGTTCT-BHQ1-3’. Viral inhibition was calculated as follow: 100* (quantity mean VC-sample quantity)/ quantity mean VC. The 50% effective concentrations (EC50 compound concentration required to inhibit viral RNA replication by 50%) were determined using logarithmic interpolation after performing a nonlinear regression (log(inhibitor) vs. response --Variable slope (four parameters)) as previously described (Touret et al., 2019, 2021a, 2022). All data obtained were analyzed using GraphPad Prism 9 software (Graphpad software).

### Viral titration

One day prior to infection, 5×10^4^ VeroE6 cells per well were seeded in 100µL assay medium (containing 2.5% FBS) in 96 well culture plates.Virus titration was performed using 96-well culture plates containing confluent VeroE6 cells inoculated with 150μL per well of four-fold dilutions of samples (dilution with medium supplemented with 2.5% FBS)Plates were incubate 6 days (37°C, 5% CO2) The absence or presence of cytopathic effect in each well was read.Infectious titers were estimated using the Reed & Muench method (REED and MUENCH, 1938).

### Mutants Generation

One day prior to infection, 5×10^5^ VeroE6/TMPRSS2 cells per well were seeded in 2mL assay medium (containing 2.5% FBS) in 12 well culture plates.The EC50s of Nirmatrelvir and Ensitrelvir were determined as previously described.Once the EC50s of these two molecules had been determined, successive passages of the virus at an MOI of 0.01 with increasing concentrations of these molecules were carried out in triplicate. Two passages at the EC50 concentration, two passages at 2x [EC50], two passages at 4 x [EC_50_] and ten passages at 8 x [EC_50_]. A no-molecule control was also run in triplicate. Between each run, the plates were incubated at 37°C/5% CO2 for 72 h. We performed extraction with Qiagen’s EZ1 automated system and NGS sequencing (Ion Torrent) for passages P8 and P16. We then performed viral production for passage P16. Viral titration by TCID_50_/mL and growth kinetics of all P16 viruses were performed.

### Growth kinetics

One day prior to infection, 5×10^6^ VeroE6/TMPRSS2 cells per well were seeded in 3mL assay medium (containing 2.5% FBS) in 6 well culture plates. The following day, the wells were infected with each virus with MOI 0.01, in 1mL of medium (containing 2.5% FBS), and incubated at 37°C/5% Co2 for 2h. Sampling at 0h took place before incubation. The experiment was performed in duplicate. After incubation, a wash with HBSS was performed and 3mL of medium (containing 2.5% FBS) was added to each well. Samples were taken at 6h pi, 24h pi, 30h pi, 48h pi, 54h pi and 72h pi. They were extracted by Qiacube as described above and qRT performed.

### Animal experiments

In vivo experiments were approved by the local ethical committee (C2EA—14) and the French “Ministère de l’Enseignement Supérieur, de la Recherche et de l’Innovation” (APAFIS#23975). All experiments were conducted in BSL 3 laboratory.

Three-week-old Syrian hamsters (Janvier labs) were maintained in ISOcage P - Bioexclusion System (Techniplast) with unlimited access to water/food and 14h/10h light/dark cycle. Animals were monitored and weighed daily throughout the duration of the study to detect the appearance of any clinical signs of illness/suffering. Groups of animals were intranasally infected under general anesthesia (isofluorane) with 50µL containing 2x10^4^ TCID50 of virus diluted in 0.9% sodium chloride solution. Animal were treated via oral route two times a day with 30 or 60 mg/kg of Ensitrelvir suspended in 0.5 % methylcellulose

Nasal washes were performed under general anesthesia (isoflurane). Blood and organs were collected immediately after euthanasia (cervical dislocation; realized under general anesthesia (isofluorane)).

Nasal washes were performed with 150µl 0.9% sodium chloride solution which was transferred into 1.5mL tubes containing 0.5mL of 0.9% sodium chloride solution, then centrifuged at 16,200g for 10 minutes and stored at -80°C. Left pulmonary lobes were washed in 10mL of 0.9% sodium chloride solution, blotted with filter paper, weighed and then transferred into 2mL tubes containing 1mL of 0.9% sodium chloride solution and 3mm glass beads. They were then crushed using a Tissue Lyser machine (Retsch MM400) for 20min at 30 cycles/s and centrifuged 10 min at 16,200g. Supernatant media were transferred into 1.5mL tubes, centrifuged 10 min at 16,200g and stored at -80°C.

Right apical lobes were collected into 2mL tubes containing 0.75mL of Qiazol lysis reagent (Qiagen) and 3mm glass beads. They were then crushed using a Tissue Lyser machine (Retsch MM400) for 10min at 30 cycles/s and stored at -80°C.

### Sequence analysis of the full-length genome

200µL of lung of infectious cell supernatant (virus stock and HAE) was inactivated with an equal volume of VXL lysis buffer (Qiagen) and viral RNA was extracted using an EZ1 Advanced XL robot with the EZ1 mini virus 2.0 kit (both from Qiagen) and linear acrylamide (ThermoFisher Scientific) in place of carrier RNA. The extracts were then subjected to quantitative real-time RT-PCR in order to standardize the amounts of viral RNA used during complete genome amplification (see below).

A specific set of primers (Table XXX) was used to generate thirteen amplicons covering the entire genome with the Superscript IV one step RT-PCR System (ThermoFisher Scientific). PCR mixes (final volume 25µL) contained 2.5µL containing a standard quantity of viral RNA from the nucleic acid extract, 0.75µL of each primer (10µM), 12.5 µL of 2X Platinum SuperFi RT-PCR Master Mix, 8.25µL of RNA free water and 0.25µL SuperScript IV RT Mix. Amplifications were performed with the following conditions: 15 sec at 55°C, 2 min at 98°C, then 40 cycles of 10 sec at 98°C, 10 sec à 56°C and 1.5 min at 72°C. Size of PCR products was verified by gel electrophoresis. For each sample, an equimolar pool of all amplicons was prepared and purified using Monarch PCR & DNA Cleanup Kit (New England Biolabs). After Qubit quantification using Qubit® dsDNA HS Assay Kit and Qubit 2.0 fluorometer (Thermo Fisher) amplicons were sonicated (Bioruptor®, Diagenode, Liège, Belgium) into 250pb long fragments. Libraries were built adding to fragmented DNA barcode for sample identification and primers with Ion Plus Fragment Library Kit using AB Library Builder System (Thermo Fisher). To pool equimolarly the barcoded samples, a real time PCR quantification step was performed using Ion Library TaqMan™ Quantitation Kit (Thermo Fisher). Next steps included an emulsion PCR of the pools and loading on 530 chips performed using the automated Ion Chef instrument (Thermo Fisher), followed by sequencing using the S5 Ion torrent technology (Thermo Fisher), following manufacturer’s instructions. Consensus sequence was obtained after trimming of reads (reads with quality score <0.99, and length <100pb were removed and the 30 first and 30 last nucleotides were removed from the reads) and mapping of the reads on a reference (MT594401.1) using CLC genomics workbench software v.21.0.5 (Qiagen). Parameters for reference-based assembly consisted of match score = 1, mismatch cost = 2, length fraction = 0.5, similarity fraction = 0.8, insertion cost = 3, and deletion cost = 3. A de novo contig was also produced to ensure that the consensus sequence was not affected by the reference sequence.

Mutation frequency for each position was calculated as the number of reads with a mutation compared to the consensus sequence divided by the total number of reads at that site.

## Public sequence data analysis

### Datasets

We downloaded all three sets of SARS-CoV-2 sequences from the GISAID public database: (1) all sequences (and attached metadata) containing an M to L change at position 49 in the NSP5 ; (2) all EG.5.1.1 sequences (filtering out low coverage and incomplete sequences) from Japan collected between July and October 2023 ; (3) all EG.5.1.1 sequences (filtering out low coverage and incomplete sequences) from North America collected between July and October 2023 ; (4) all sequences (and attached metadata) containing an M to I change at position 49 in the NSP5 .

We first removed all sequences with >5% of ambiguous nucleotides from all datasets except for the one corresponding to EG.5.1.1 M49L sequence. Then, to obtain a computationally tractable dataset for phylogenetic inference, we randomly downsampled the EG.5.1.1 North America dataset (filtering out duplicates) to 500 sequences, and combined it with all EG.5.1.1 sequences from Japan. We added all EG.5.1.1 sequences carrying the M49L mutation. After the first maximum likelihood (ML) inference, we removed three sequences corresponding to extremely long terminal branches (GISAID Accession IDs: 18434659, 18226914, 18432517), yielding a final dataset of 2603 sequences. All GISAID accession numbers are available in Supp. Table 2, the alignment and trees are available at https://github.com/rklitting/SARS-CoV-2-resistance.

### Phylogenetic analysis

For the M49L and M49I sequences lineage analysis, we assigned lineages to each of the M49L sequences using the Pangolin lineage assigner (version 4.3)(O’Toole et al., 2021). Classification failed for 3 out of the 267 sequences submitted (See Supp. Table 1).

For phylogenetic inference based on M49L-sequences, we aligned the final EG.5.1.1 dataset to the reference genome WIV04 (GISAID Accession: 402124) using MAFFTv7(Katoh and Standley, 2013). To provide some phylogenetic context to the EG.5.1.1 lineage, we added the set of genomes corresponding to WHO reference strains for VOCs, VUIs and VUMs provided by the Los Alamos national laboratory(Korber et al., 2020) .Finally, we performed the masking of sites previously identified as potential sequencing errors or suspect homoplasies(“Issues with SARS-CoV-2 sequencing data - SARS-CoV-2 coronavirus / nCoV-2019 Genomic Epidemiology,” 2020), and removed the 5’ and 3’ UTRs. We inferred a first ML phylogeny for this dataset using IQTREE2 under ModelFinder with ultrafast bootstrap approximation (1000 replicates)(Minh et al., 2020, 2013). Finally, we performed 10 ML inference replicates, to control for potential instability in the topology, in particular within the EG.5.1.1 lineage.

### Visualizations

The plots were created using matplotlibv3.6.3(Hunter, 2007)and the tree was visualized using baltic (https://github.com/evogytis/baltic).

## Results

### Generation of SARS-CoV-2 strains resistant to protease inhibitors

To generate mutants resistant to SARS-CoV-2 protease inhibitors, we first determined the experimental conditions suitable for the emergence of such resistance mutations in cell culture. We evaluated, for both molecules, the half maximal effective concentration EC_50_ in the VeroE6 TMPRSS2 cell line, using remdesivir as a reference compound and the BavPat1 SARS-CoV-2 strain (B.1 lineage carrying the D614G mutation). In line with the literature(Imai et al., 2022; Sasaki et al., 2022; Takashita et al., 2022; Unoh et al., 2022), we obtained EC_50s_ of 0.14µM and 2.26µM for ensitrelvir and nirmatrelvir, respectively (Fig1a). The EC_50_ of nirmatrelvir appears to be higher than that of ensitrelvir, as we chose to use similar conditions for all our *in vitro* experiments and therefore did not add an efflux pump inhibitor with nirmatrelvir, as other groups may have done (Boras et al., 2021; Owen et al., 2021).

**Figure 1:**
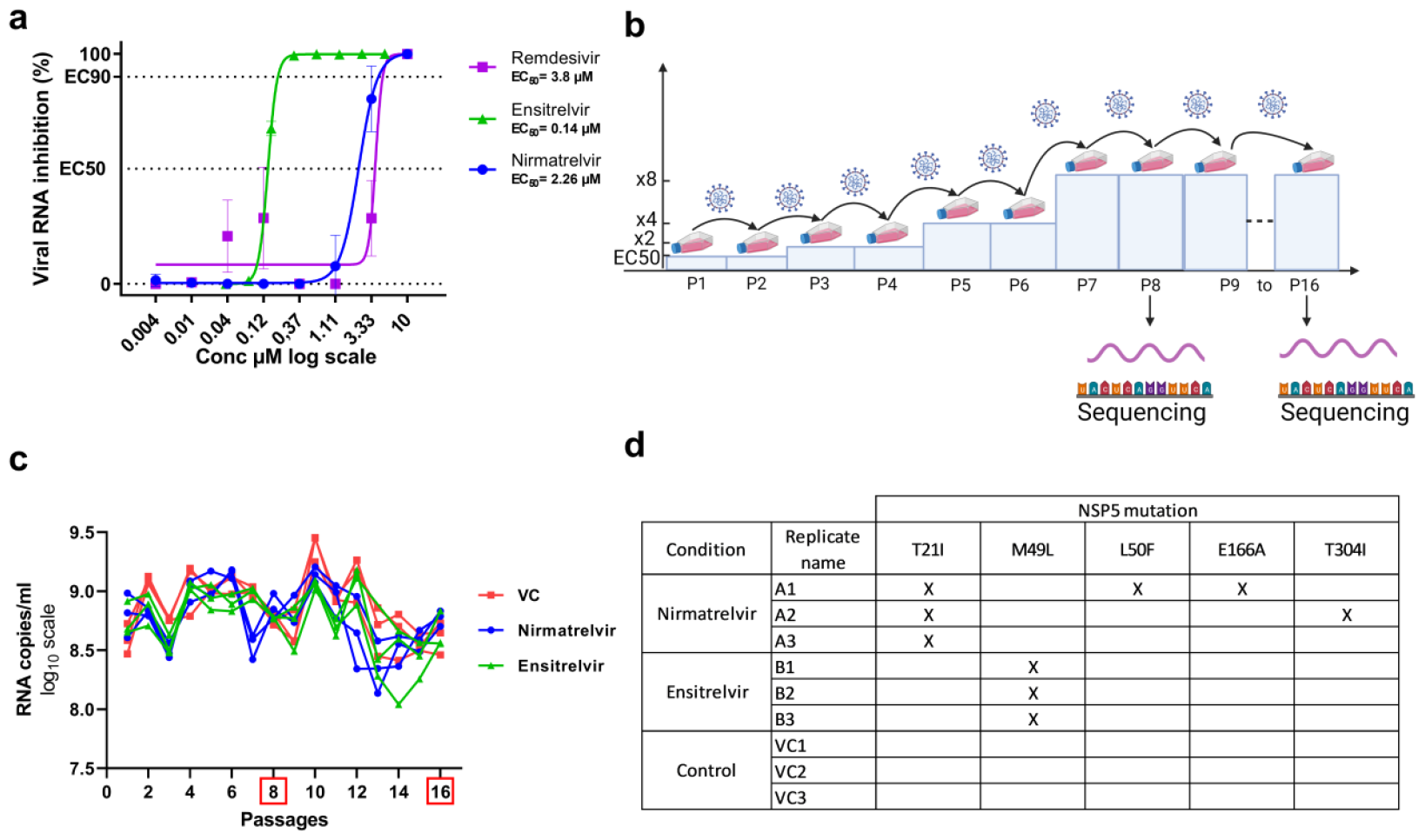
Generation of nirmatrelvir and ensitrelvir resistant strains. a) Pilot experiment to determine the EC50s of nirmatrelvir and ensitrelvir. b) Graphical representation of the strategy used to obtain resistant strains. c) Viral replication measured by the amount of viral RNA at each passage in each replicate. d) Table of NSP5 amino acid mutations obtained at passage 16 in the different replicates

Then, for each of the compounds, we started to grow the virus at the EC_50_, doubling the drug amount every two passages to increase selective pressure while allowing the virus to replicate (Fig1b). In parallel, we performed virus passages in the absence of antiviral molecules to identify cell line adaptation mutations. We performed the experiment with 3 replicates of each condition. To assess the emergence of potential resistance mutations at passage 8, we sequenced the NSP5 gene -which contains the 3CLpro region-but found no mutations. As it was not possible to further increase the concentration of antiviral molecule in the supernatant, we carried out 8 additional passages at the maximum concentration (8 times higher than the EC_50_). Throughout the experiment, we were able to maintain a stable level of viral replication without any remarkable drop in viral load (Fig. 1c).

By sequencing full viral genomes from all experimental conditions at passage 16, we identified mutations in the 3CLpro region, including some that were conserved across all replicates (Fig1d).

All three replicates of the cultures performed in the presence of nirmatrelvir exhibited the T21I mutation. The other mutations -T304I, L50F, E166A-were observed in one out of three replicates. We observed T21I either alone (replicate A3), combined with T304I (A2), or combined with both L50F and E166A (A1). All four mutations have been previously described as conferring nimatrelvir resistance(Hu et al., 2023; Iketani et al., 2023; Kawashima et al., 2023).

In the case of ensitrelvir, we only observed a single mutation, M49L, conserved across the three replicates. This mutation has already been described both in clinical isolates(Ip et al., 2023; Kawashima et al., 2023) and following experiments conducted *in vitro* (Kiso et al., 2023b) and shown to reduce sensitivity to the drug.

### *In vitro* characterization of resistant strains

To characterize our mutant strains *in vitro*, we first monitored viral replication in VeroE6 TMPRSS2 cells in the absence of compound (Fig.2a), using a low MOI (0.02), and a RT-qPCR assay.

Based on molecular viral load, we found no significant difference among the strains tested. These results are in line with the literature for T21I, T21I+T304I (Iketani et al., 2023) and M49L(Kiso et al., 2023b). They indicate that the mutations emerged in cell culture in the presence of nirmatrelvir and ensitrelvir do not alter significantly *in vitro* replication (Fig.2a)

Using a dose-response approach, we then characterized the sensitivity of these strains to the two protease inhibitors, using remdesivir as a control. In addition to our wild-type reference strain (BavPat), we tested one of the strains that had undergone the same number of passages as the resistance mutants but without inhibitors (VC3). For the remdesivir control, we obtained very similar EC_50_ values for the 8 viruses tested, with a maximum variation of 0.5µM in their EC_50_ (Fig 2d, Table 1). This indicates that the 16 passages in culture did not alter the sensitivity of the strains to remdesivir or their basic replication properties.

**Figure 2:**
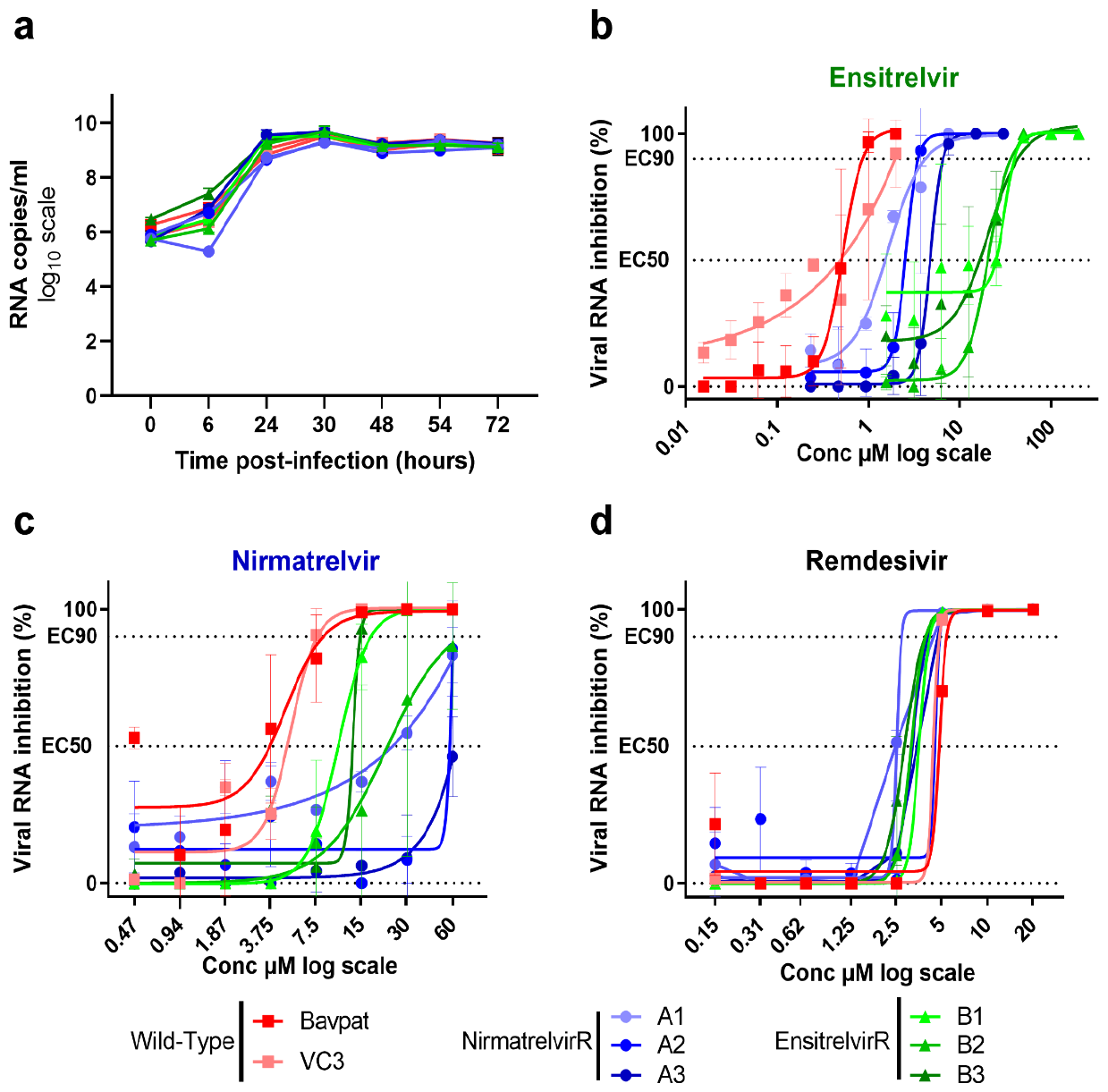
In vitro characterization of nirmatrelvir and ensitrelvir resistant strains. a) Kinetics of replication in VeroE6 TMPRSS2 cells Data presented are from two technical replicates and error bars show mean±s.d ; Dose response curves reporting the susceptibility of SARS-CoV-2 mutants and two wild type strain against b) ensitrelvir c) nirmatrelvir and d) remdesivir. Data presented are from three technical replicates in VeroE6-TMPRSS2 cells, and error bars show mean±s.d

**Table 1:**
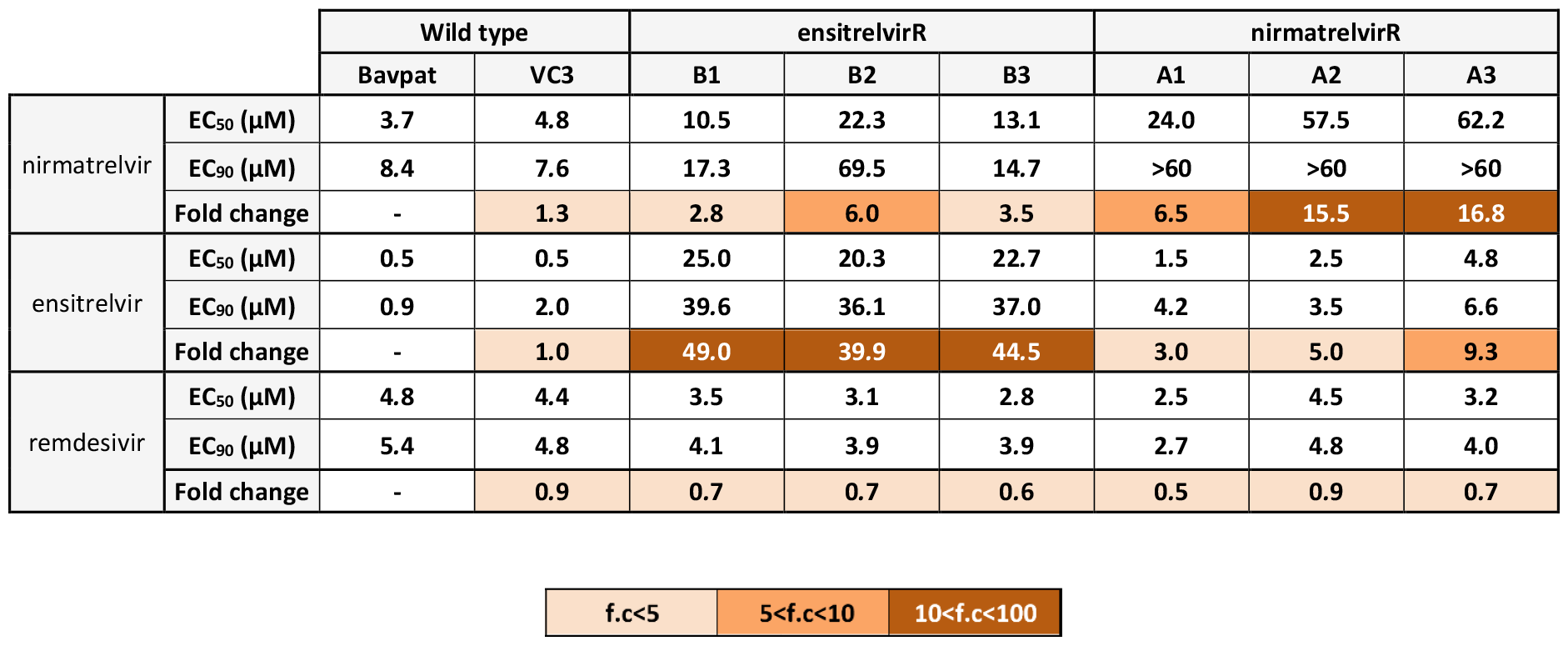
Activity of ensitrelvir, nirmatrelvir and remdesivir against wild type and mutants strains. Interpolated EC_50_ values are expressed in µM. Fold change reductions were calculated in comparison with the EC_50_ of the BavPat strain. F.c : fold change

|  |  | Wild type |  | ensitrelvirR |  |  | nirmatrelvirR |  |  |
| --- | --- | --- | --- | --- | --- | --- | --- | --- | --- |
|  |  | Bavpat | VC3 | B1 | B2 | B3 | A1 | A2 | A3 |
| nirmatrelvir | EC<sub>50</sub> (μM) | 3.7 | 4.8 | 10.5 | 22.3 | 13.1 | 24.0 | 57.5 | 62.2 |
|  | EC<sub>90</sub> (μM) | 8.4 | 7.6 | 17.3 | 69.5 | 14.7 | >60 | >60 | >60 |
|  | Fold change | - | 1.3 | 2.8 | 6.0 | 3.5 | 6.5 | 15.5 | 16.8 |
| ensitrelvir | EC<sub>50</sub> (μM) | 0.5 | 0.5 | 25.0 | 20.3 | 22.7 | 1.5 | 2.5 | 4.8 |
|  | EC<sub>90</sub> (μM) | 0.9 | 2.0 | 39.6 | 36.1 | 37.0 | 4.2 | 3.5 | 6.6 |
|  | Fold change | - | 1.0 | 49.0 | 39.9 | 44.5 | 3.0 | 5.0 | 9.3 |
| remdesivir | EC<sub>50</sub> (μM) | 4.8 | 4.4 | 3.5 | 3.1 | 2.8 | 2.5 | 4.5 | 3.2 |
|  | EC<sub>90</sub> (μM) | 5.4 | 4.8 | 4.1 | 3.9 | 3.9 | 2.7 | 4.8 | 4.0 |
|  | Fold change | - | 0.9 | 0.7 | 0.7 | 0.6 | 0.5 | 0.9 | 0.7 |

|  |  |  |
| --- | --- | --- |
| f.c<5 | 5<f.c<10 | 10<f.c<100 |

**In the ensitrelvir sensitivity analysis**, the VC3 strain at 16 passages showed a sensitivity very close to that of the BavPat strain, with an EC_50_ of 0.49µM versus 0.51µM. This rules out an effect of the number of passages on sensitivity (Fig. 2b, Table 1). Strains selected in the presence of nirmatrelvir (nirmatrelvirR) showed a slightly reduced sensitivity to ensitrelvir (3 to 9-fold). On the other hand, the three M49L ensitrelvir-resistant (ensitrelvirR) mutants showed a great decrease in sensitivity (around 40-fold), indicating that this mutation does confer resistance to ensitrelvir (Fig. 2b, Table 1).

**In the nirmatrelvir sensitivity analysis**, the VC3 strain showed a sensitivity close to that of the BavPat strain, with an EC_50_ of 4.82µM versus 3.70µM (Fig. 2cTable 1). As before, two profiles were observed for the mutant strains (Fig. 2cTable 1): ensitrelvirR strains showed a slightly reduced sensitivity to nirmatrelvir (3 to 6-fold). On the other hand, the three nirmatrelvirR mutants showed a greater decreased in sensitivity (6 to 17-fold) (Fig. 2c Table 1). Interestingly, strains nirmatrelvirR-A2 and nirmatrelvirR-A3 showed the greatest resistance, despite having only one and two mutations, respectively. These results confirm that the T21I mutation and the combinations T21I-T304I and T21I-L50F-E166A do confer resistance to nirmatrelvir.

### Ensitrelvir is not efficacious against M49L ensitrelvir -resistant strain *in vivo*

Nirmatrelvir mutants have already been extensively studied(Heilmann et al., 2022; Hu et al., 2023; Iketani et al., 2023; Kiso et al., 2023a) but the ensitrelvir resistance mutation M49L has never been evaluated alone *in vivo* in an antiviral susceptibility assay. To assess the ability of this mutation to confer resistance to ensitrelvir *in vivo*, we used a hamster model of SARS-CoV-2 infection previously developed for antiviral evaluation(Driouich et al., 2021; Touret et al., 2021b; Driouich et al., 2022). We infected groups of 6 hamsters intranasally with either the wild type virus (BavPat), or M49L-R strain (B2). In addition to untreated groups, hamsters were administered 30 or 60 mg/kg of ensitrelvir orally twice a day. This regimen has previously been validated in two murine models(Kiso et al., 2023b; Unoh et al., 2022). It should be noted that in our study, unlike the one exploring loss of sensitivity to ensitrelvir in vivo, we used the strain from repeated passages and not generated by reverse genetics.

We observed that the M49L-ensitrlevirR strain and the wild type exhibited similar levels of replication in lungs of untreated hamster (no statistical differences) both in terms of viral RNA yield and infectious titers (Fig. 3 a and b). Regarding clinical follow-up, both the M49L-R strain and the wild-type induced a similar weight loss in hamsters (Fig. 3 c and d). These results show that, as observed *in vitro*, the selection process for resistance mutations has not deeply altered the replication characteristics and the pathogenicity of this strain in our model.

**Figure 3:**
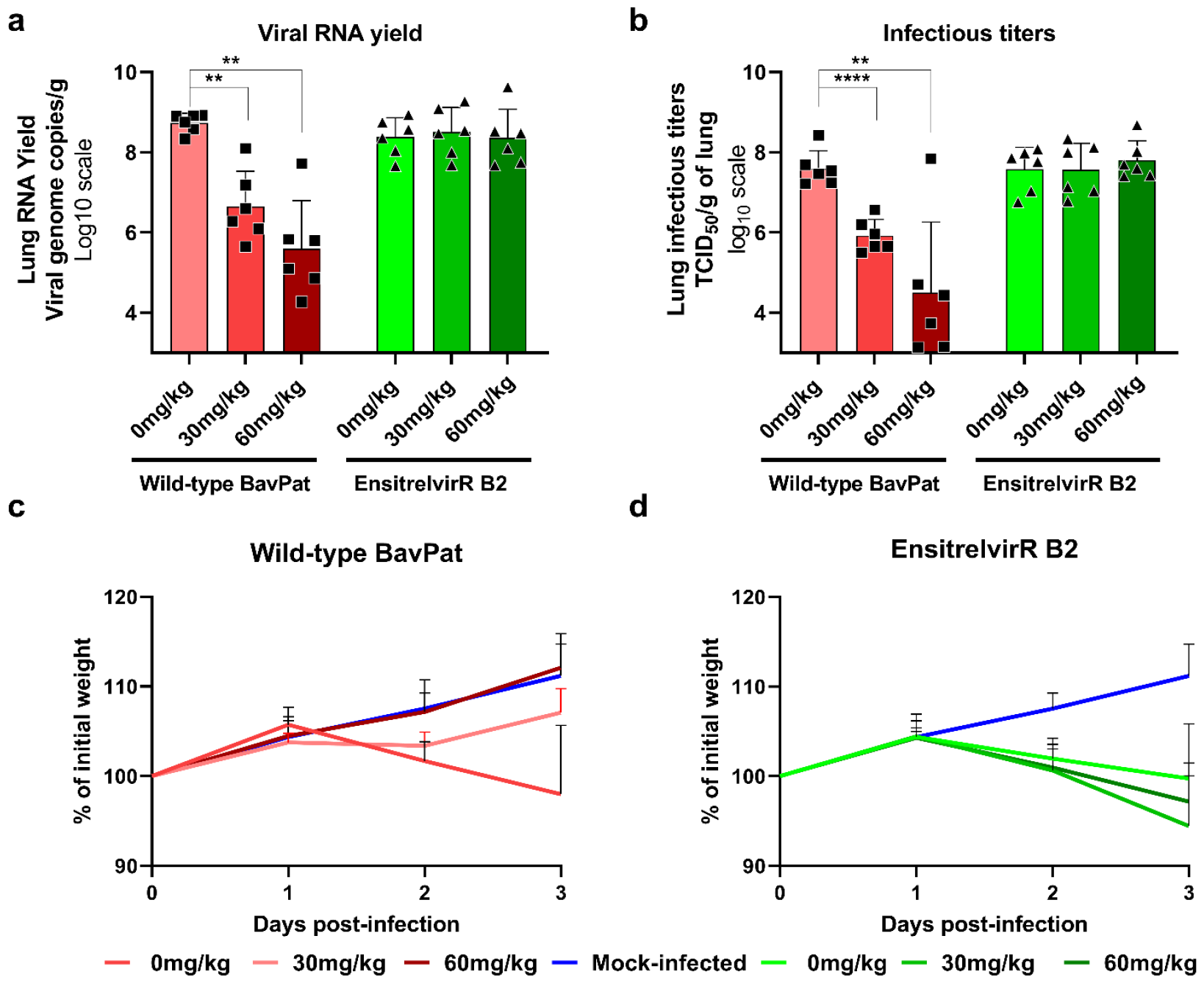
Antiviral activity of oral ensitrelvir treatment in a hamster model against wild type and resistant SARS-CoV-2 strains. Groups of 6 hamsters were intranasally infected with 10^4 TCID50 of virus. Animals received ensitrelvir orally twice a day at 0 (untreated group), 30 or 60 mg/kg. a) Viral replication in lung based on viral RNA yields at day 3 post infection (measured using an RT-qPCR assay) expressed in viral genome copies/g of lung. b) Viral replication in lung based on infectious titers at day 3 post-infection (measured using a TCID50 assay) expressed in TCID50/g of lung. c-d) Clinical course of the disease (n=6 animals/group). Normalized weight at day n was calculated as follows: % of initial weight of the animal at day n. Data represent mean±SD. Two-sided statistical analysis was performed using Shapiro-Wilk normality test and Student t-test, ** and **** indicate and average significant value lower than that of the untreated group, with a p-value ranging between 0.001-0.01 and lower to 0.0001 respectively.

**Regarding the antiviral effect of ensitrelvir on the wild-type strain**, we observed a significant decrease in the amount of viral RNA and infectious titer in lungs, with a dose-response effect (Fig 3 a and b). Furthermore, animals treated with 60 mg/kg had a similar clinical follow-up to uninfected hamsters with no statistical differences in weight change, suggesting a beneficial effect of treatment on clinical outcome (Fig 3 c and Supplemental Fig.1). Animals treated with 30 mg/kg had an intermediate clinical follow-up between untreated infected hamsters and uninfected hamsters, with a similar trend to uninfected hamsters despite a decline between day 1 and 2 (Fig. 3 c and Supplemental Fig.1). These results indicate significant antiviral activity of ensitrelvir at both doses tested on the wild-type strain.

**Regarding the antiviral effect of ensitrelvir on the resistant strain**, we observed no significant decrease in viral RNA levels or infectious titers in lungs compared to control, regardless of the dosing regimen (Fig. 3 a and b). Furthermore, animals treated with 30 and 60 mg/kg had a clinical follow-up similar to untreated infected hamsters with no statistical differences in weight change to untreated infected hamsters, suggesting no beneficial effect of treatment on clinical outcome (Fig. 3d and Supplemental Fig.1). These results indicate a lack of efficacy of ensitrelvir at both doses tested on the mutant strain containing the M49L mutation in NSP5. This result is in line with that obtained with the M49L/E166A double mutant in the same hamster model(Kiso et al., 2023b), and demonstrates the ability of M49L to confer resistance to ensitrelvir even if in the absence of E166A.

### The M49L mutation increased in prevalence over the last 6 months

The emergence of resistance mutations can drastically limit the use of the antiviral therapy affected. Resistance may emerge during prolonged use of the molecule in a patient with ongoing viral replication but can also occur naturally – without any specific selective pressure due to the use of the molecule in question. Our experiments show that the M49L mutation alone can confer resistance to ensitrelvir and previous reports have described M49L in circulating SARS-CoV-2 variants(Ip et al., 2023; Kawashima et al., 2023). The prevalence of this mutation in recent months and in particular, since the commercialization of ensitrelvir in Japan, has, however, not been reported.

We therefore searched on GISAID for all SARS-CoV-2 sequences carrying the M49L mutation. We found that, as of 30^th^ of October 2023, only 267 (0.00165%) of the 16,167,749 SARS-CoV-2 sequences available on GISAID present an M to L amino acid substitution at position 49 in the NSP5 gene. Using sequence collection dates, we evaluated the distribution over time of all M49L-carrying SARS-CoV-2 sequences. We found that M49L was first reported on the 13^th^ August 2020, and has been observed almost monthly since then (Figure 4a), with an apparent global prevalence of 0.00168% (total number of sequences with collection date on or after 13^th^ August 2020: 15,929,475). When looking at the most recent time span, we found that more M49L-carrying sequences have been reported between May and October 2023 (count: 154; prevalence: 0.036%) than over the previous 21 months (count: 113 M49L; prevalence: 0.0007%). These numbers indicate that while M49L corresponds to a very small proportion of the sequences observed until today, its prevalence has increased during the last 6 months.

**Figure 4:**
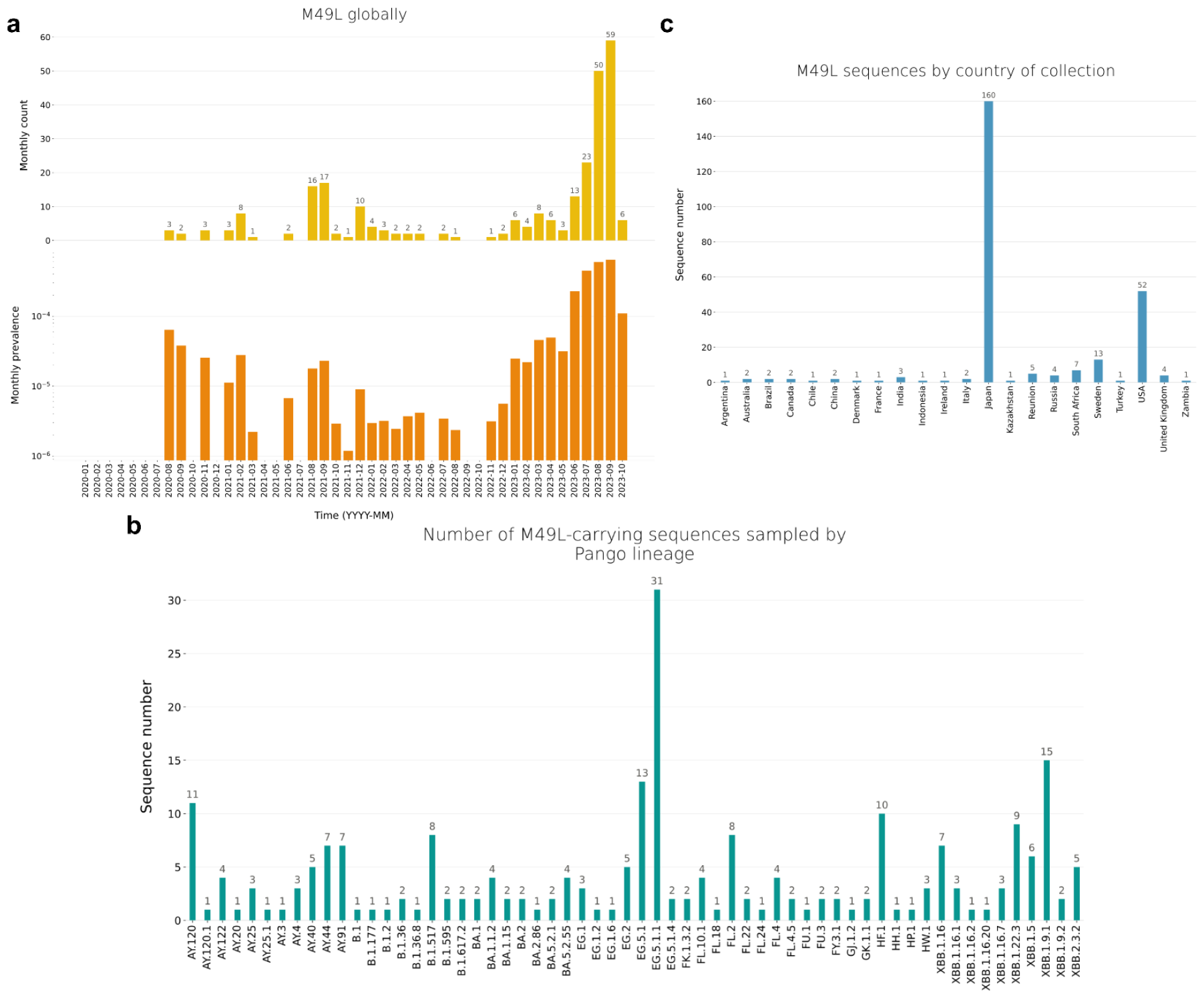
M49L distribution in time, space, and across lineages. **a)** Global monthly counts (upper panel) and prevalences (lower panel) of M49L-sequences, based on all SARS-CoV-2 genomes publicly available on GISAID as of 2023-11-13. b) Number of M49L sequences observed by lineage, for each of the 60 Pango lineages observed most frequently among M49L sequences. c) Number of M49L sequences observed by country of collection.

The repeated observations of low numbers of M49L-carrying sequences since the first year of the pandemic suggest that the mutation has emerged on multiple occasions and in several SARS-CoV-2 lineages. To determine the Pango lineage of each of the 267 sequences with the M49L mutation, we used the Pangolin online lineage assigner(O’Toole et al., 2021). We found that M49L-carrying sequences are distributed across 90 different lineages, including Alpha, Delta, and Omicron, with up to 31 sequences belonging to the same lineage (Figure 4b, Supp. Table 1). The most prevalent lineages among M49L-carrying sequences are EG.5.1.1 (11.6%), XBB.1.9.1 (5.6%), and EG.5.1 (4.9%). These three lineages are “active” (last observed less than a year ago) and classified either as “variant of interest” (VOI) or as “variant under monitoring” (VUM) by the WHO(“Tracking SARS-CoV-2 variants,” n.d.). Overall, these results indicate that NSP5 M49L can emerge in multiple genomic backgrounds and is present in active, currently prominent, lineages.

### The recent increase in the prevalence of M49L appears to be linked to a specific selective pressure in Japan

Resistance against an antiviral therapy may occur naturally in circulating lineages but the use of a drug may favor the emergence of resistance mutations. Using the collection dates of M49L-carrying sequences, we showed that their proportion has been multiplied by more than 50 times in the last 6 months. Several scenarios may explain this increase in observation frequency: (i) a specific selective pressure; (ii) a more favorable genomic background; (iii) increased transmission.

The increased prevalence of the M49L resistance mutation observed in this study may be due to the existence of specific selective pressure, *i*.*e*. the use of the drug to which the compound confers resistance, ensitrelvir. The only country where this drug is used is Japan, where it first received an emergency use approval in November 2022, and was made commercially available in April 2023(“Xocova to Be Commercially Available from March 31,” n.d.). When we evaluated the spatial distribution -in terms of country of origin-of M49L sequences, we found that Japan is by far the most important country of origin for M49L sequences, with 59.9% of sequences originating in Japan overall, a number that increases to 88.3% when considering only collection dates between May and October 2023 (Figures 4c and 5a). These results indicate that the vast majority of the M49L-carrying sequences collected over recent months originate from Japan.

When analyzing the time distribution of all M49L-sequences from Japan, we found that M49L has been observed only 4 times before December 2022 but almost monthly since then (Figure 5b), with especially high monthly prevalence values (above 0.2%) since June 2013. The timing of the increase in the prevalence of M49L and the origin of M49L sequences both align well with the use of Xocova® in Japan, suggesting that this selective pressure may favor the emergence of the mutation.

**Figure 5:**
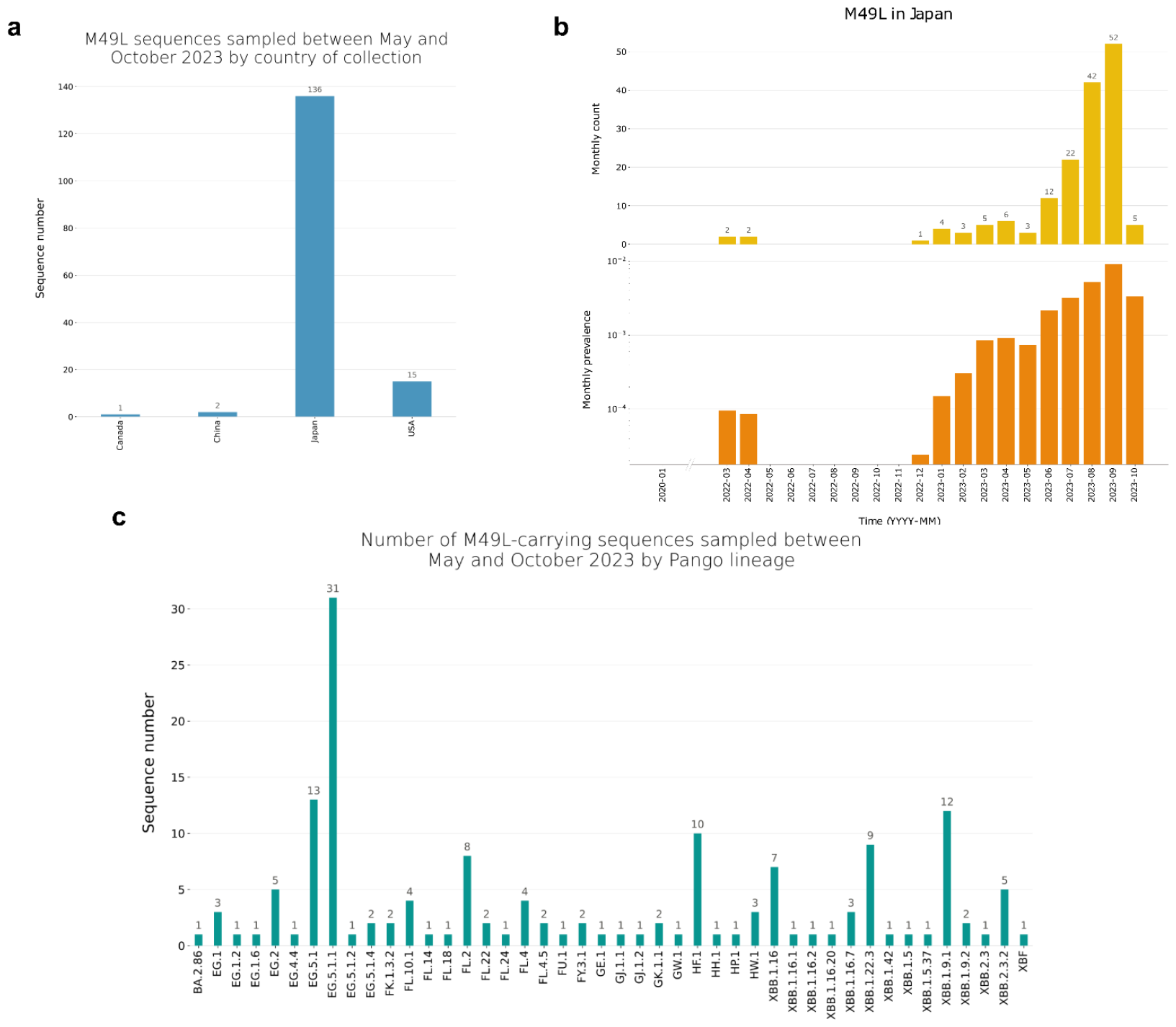
M49L distribution in time, space and, across lineages between May and October 2023 in Japan. **a)** Number of M49L sequences observed by country of collection for sequences collected between May and October 2023. b) Monthly counts (upper panel) and prevalences (lower panel) of M49L-sequences in Japan, based on all SARS-CoV-2 genomes publicly available on GISAID as of 2023-11-13.c) Number of recent M49L sequences (collection date between October and May 2023) observed by lineage, for each of the Pango lineages observed among recent M49L sequences.

Another factor that may however, explain the recent increase in M49L sequences overall and in Japan more specifically, is the local circulation of lineages with a genomic background more favorable for the emergence of the mutation. To evaluate if recent M49L-sequences were associated with a specific genomic background, we analyzed their lineage distribution. We found that between May and October 2023, M49L occurred in 44 different lineages globally and in 40 different lineages in Japan including, EG.*, FL.*, GJ.*, HF.*, XBB.1.9.*, and XBB.1.16.* lineages (Figure 5c). These results indicate that multiple distinct lineages can accommodate the M49L mutation.

When considering M49L-sequences from the most prevalent lineage, EG.5.1.1, 93.5% of them appear to originate in Japan, and only 3.2% both in the US and in Canada. Based on GISAID data, EG.5.1.1 appears to have circulated to levels on the same order of magnitude (or greater) to that observed over the last 6 months in Japan (7.6% of 37550 total sequences), in the US (3.8% of 146709 total sequences), in Canada (6.8% of 46555 total sequences), and in China (18.2% of 34404 total sequences). If the increase in M49L prevalence was driven by the genomic background of EG.5.1.1 being more favorable, this mutation would have also emerged more frequently in other locations where the lineage has established sustained transmission. These observations indicate that the recent increase in M49L prevalence is not driven by a single -or a few-more favorable genomic background.

### Repeated emergence, rather than increased transmission, contributed to the recent increase in M49L prevalence

Another phenomenon that may have participated in increasing the prevalence of M49L in recent month is the formation of large transmission chains by naturally-occurring M49L mutant. The distribution of recent M49L sequences across 44 different lineages suggests that increased emergence events, rather than increased transmission, fueled the rise in M49L detections. However, the presence of a large transmission chain within the main lineage EG.5.1.1 may have been an exacerbating factor.

To determine whether recent M49L-carrying sequences from the most prominent lineage, EG.5.1.1, correspond to a single or multiple emergence events, we performed a phylogenetic inference based on a set of publicly available SARS-CoV-2 genomes. Using a maximum-likelihood approach, we inferred the phylogenetic relationships between all M49L-carrying EG.5.1.1 sequences and a set of EG.5.1.1 sequences collected between July and October in Japan and North America (to match the spatio-temporal distribution of the M49L genomes). In the resulting phylogeny, we identified 12 different M49L emergence events, associated either with single tips, or with entire well-supported clades encompassing up to 10 sequences (Figure 6). Our findings were broadly similar across 10 inference replicates. These results show that recent M49L-carrying sequences do not form a single clade -- the mutation emerged at least on 12 independent occasions within that lineage.

**Figure 6:**
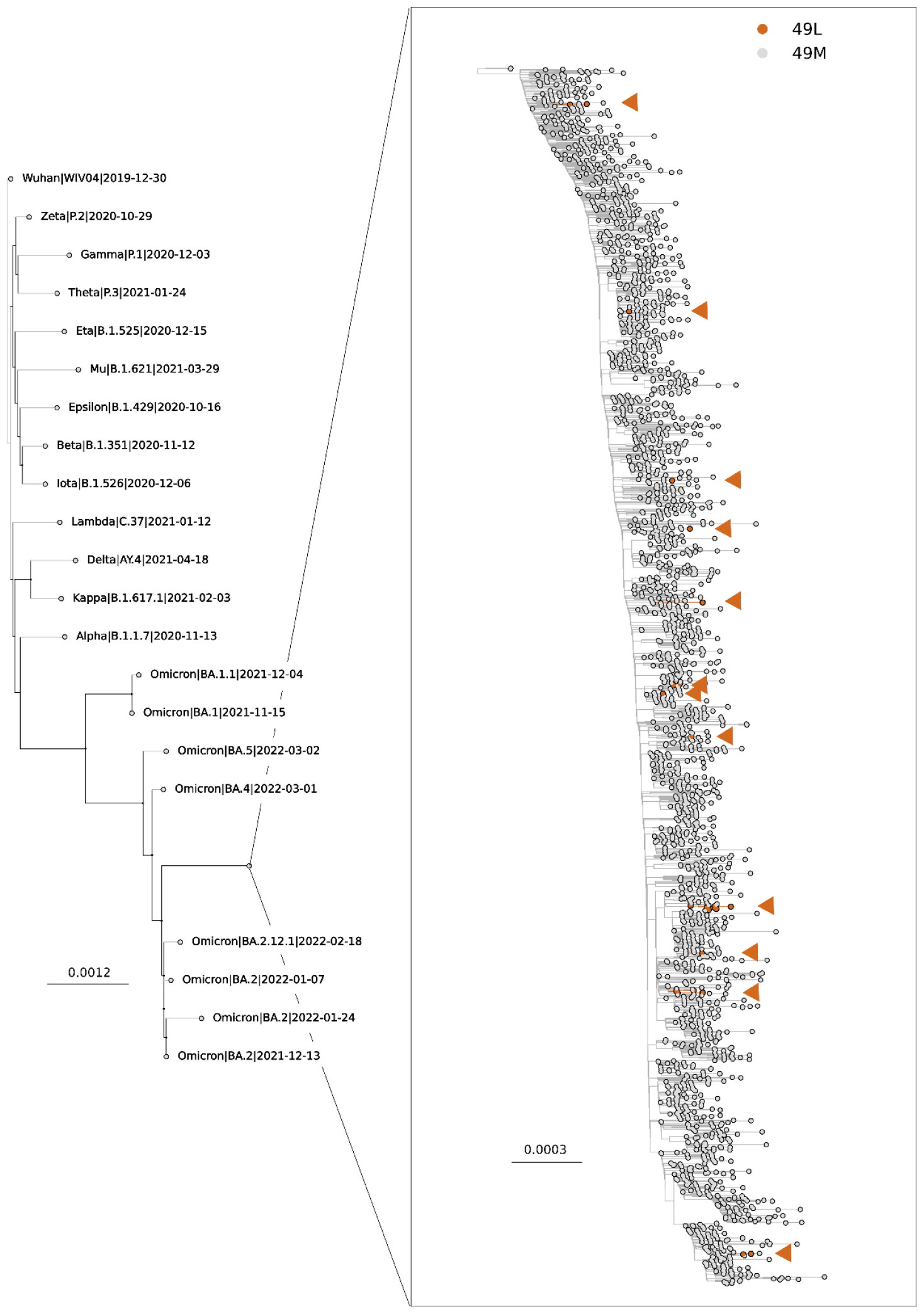
Maximum-likelihood phylogeny with a zoom on the EG.5.1.1 lineage. Branches and tips associated with M49L-carrying sequences are highlighted in orange. Orange arrows indicate M49L emergence events.

## Discussion

In this study, we experimentally generated six viruses carrying resistance mutations to clinical stage SARS-CoV-2 protease inhibitors. We obtained three viruses with different sets of mutations for nirmatrelvir. For ensitrelvir, on the other hand, all three strains had the same mutation M49L. *In vitro* characterisation showed that nirmatrelvir-resistant strains had low resistance to ensitrelvir, but higher resistance to nirmatrelvir. Similarly, ensitrelvir-resistant strains showed low resistance to nirmatrelvir but much higher resistance to ensitrelvir. Finally, we investigated *in vivo* the resistance of an EnsitrelvirR M49L strain to two regimens of ensitrelvir and found that it completely escaped the inhibition induced by this antiviral compound.

We obtained resistance mutations after 16 passages with an increasing concentration of nirmatrelvir, which is similar to what has already been shown by Iketani and colleagues(Iketani et al., 2023). Indeed, these specific mutations have already been obtained using a similar approach: the T21I mutation was generated alone or in combination with E166V or T304I, exactly as in our study. This mutation, along with T304I, was predominantly acquired as initial mutation(Iketani et al., 2023). Interestingly, in the VeroE6 TMPRSS2 cell line, we obtained a mutation at residue 166, whereas in the study by Iketani and colleagues(Iketani et al., 2023) a mutation at this position only appeared in HUH.7 and not in VeroE6, and the authors supposed that it was specific to this cell line. The mutation on residue 166 had already been described and was shown to reduce viral replicative fitness, but this fitness could be restored by the addition of the T21I mutation(Iketani et al., 2023). In our study, we confirmed this observation by showing that the replication capacity of our mutant (A1) carrying both T21I and E166A was the same as that of the wild type. Finally the mutation at residue 166 was the only one obtained in the substrate binding site; the others are situated at a distal position from nirmatrelvir predicted binding site (over 5 Å(Iketani et al., 2023)).

For ensitrelvir, we obtained three viruses with the same mutation, M49L. This mutation has already been obtained in a similar experiment, but with a shorter passage number(Kiso et al., 2023a). It is also present in circulating isolates of SARS-CoV-2 (Hu et al., 2023; Kawashima et al., 2023; Moghadasi et al., 2023b; Noske et al., 2023).

In a report identifying M49I as an ensitrelvir resistance mutation its visualization in complex with the 3CLpro shows that the greater hydrophobicity of isoleucine, compared to the methionine, caused the inhibitor to move to another site, which affects its antiviral activity(Noske et al., 2023). According to another study, the presence of a leucine instead of an isoleucine at this position could cause a greater resistance due the gamma-carbon branching of the leucine side chain, which is nearer to H41 one of the catalytic dyad in its active site (Moghadasi et al., 2023b; Unoh et al., 2022).

We found *in vitro* moderate cross resistance between NirmatrelvirR strains and ensitrelvir, and the same for EnsitrelvirR strains and Nirmatrelvir, in accordance with previous studies(Iketani et al., 2023; Moghadasi et al., 2023b). This suggests that replacing one of the two compounds with the other could compensate for the emergence of a resistance mutation during treatment.

We observed the greatest loss in activity with the couple ensitrelvir/M49L with a ~40-fold reduction in susceptibility for our three ensitrelvirR strains. This mutation had already been tested, alone, in two distinct enzymatic assays(Moghadasi et al., 2023b, 2023a) and in a VSV based cleavage system(Moghadasi et al., 2023b), with observed fold change values of 21.4, 25.4 and 10 respectively. In the only previous study that explored the effect of this mutation with a replicative virus (a mutated Delta strain generated by reverse genetic(Kiso et al., 2023b)) M49L was associated with a 37.4 fold change in a focus reduction assay(Kiso et al., 2023a). This result is perfectly in line with our observation with the same range of reduction despite the use of the strain generated from repeated passages in a different variant backbone. The fact that we found the same effect for a single amino acid mutation but with a different backbone not only confirms but reinforces our result.

In the abovementioned study, the authors only studied the *in vivo* antiviral resistance of the M49L/166A mutation pair(Kiso et al., 2023b). Regarding the single mutation M49L, they only explored its impact on the *in vitro* replication fitness and the *in vivo* pathogenicity. We went a step further and explored the effect of this single mutation on ensitrelvir sensitivity, using both the ensitrelvir dose referred to in the study above (60mg/kg twice daily)(Kiso et al., 2023b) and a lower dose as in the original S-217622/ensitrelvir article (30mg/kg twice daily)(Unoh et al., 2022). When animals were infected with the ensitrelvirR strain, we observed no antiviral activity for either of ensitrelvir dose regimen and, in contrast to what was observed with the wild-type strain, no clinical improvement. This implies that the M49L mutation can be solely responsible for *in vivo* resistance. The same result at the highest regimen was observed *in vivo* with the double mutants M49L/E166A(Kiso et al., 2023b). The *in vitro* fold change of this combination is almost 200(Kiso et al., 2023b). Our result implies that a single mutation inducing a change in susceptibility of 40, *i*.*e*. five times lower, is enough to make ensitrelvir treatment ineffective in our animal model.

In the current study, we detailed a recent increase in the prevalence of M49L-carrying sequences, linked it to a specific region, Japan, and showed that this increase is concomitant with the use of ensitrelvir in the country (emergency approval and then commercialization) rather than with the emergence of a more favorable viral genomic background. These initial results are concerning and warrant further investigation, especially as several countries are currently considering the drug for emergency approval(“Drugmaker Shionogi inks deal to market COVID pill in China,” n.d.; “Xocova Filed for Conditional Approval in South Korea,” n.d.). It is crucial to better characterize the conditions and frequency of emergence of the M49L mutation in patients treated with ensitrelvir. If an increased risk of resistance emergence during patient treatment is confirmed, genomic monitoring may allow to both optimize the medical efficacy of the treatment provided and prevent the spread of resistant strains. In case of resistance, it would still be possible to replace ensitrelvir with nirmatrelvir, given the low level of cross-resistance between the two molecules. However, this would be difficult for patients who had received ensitrelvir treatment in the first instance, due to a contraindication to ritonavir, which is present in Paxlovid®, the pharmaceutical formulation of nirmatrelvir.

To evaluate the potential duration of transmission for EG.5.1.1 M49L-mutants, we calculated minimum clade-durations for all M49L-clades by calculating the difference in time between the earliest and latest sampling dates among sequences of a given clade. We found that clade-duration could be up to 58 days for M49L clades belonging to the EG.5.1.1 lineage. This first assessment however, is based on a small number of clades (3) and on a limited sampling of the recent circulating diversity of SARS-CoV-2. For computational tractability reasons we focused our phylogenetic analysis on sequences from a single lineage (EG.5.1.1), specific countries (Japan, North America), and subsampled our dataset. We believe that it is of the utmost importance to pursue these investigations and to continue to monitor the ability of M49L-carrying lineages to establish sustained transmission and to assess the risk of fixation of the mutation.

Finally, while we showed that the use of ensitrelvir is probably a driver of the increased frequency of emergence of M49L in recent months in Japan, it is likely that the 107 early sequences carrying M49L (identified between August 2020 and October 2023) emerged spontaneously as the antiviral was not in use before November 2022. The natural occurrence of strains carrying M49L may thus facilitate the emergence of resistance to ensitrelvir.

Other variations in the NSP5 protein provide resistance to ensitrelvir and have been observed in circulating isolates. They may present a risk of increased emergence in case of wide use of ensitrelvir, akin to what was showed for M49L in the present study. In particular, the mutation M49I provides resistance *in vitro* (Moghadasi et al., 2023b) and is present in 2188 SARS-CoV-2 sequences on GISAID (as of October 30th). For this mutation however, we found no apparent increase in prevalence over recent months in Japan (Supplemental Figure 2a) with most M49I sequences observed between June and December 2021. Along with the fact that M49I has never been observed in experiments of generation of ensitrelvir-resistant mutants (current study and in (Kiso et al., 2023b), this finding suggests that under selective pressure, the M49L mutation is preferred, likely because it provides greater resistance by generating a branching pattern more favorable to the disruption of a base stacking interaction with the inhibitor (Moghadasi et al., 2023b; Noske et al., 2023). In addition, the lower prevalence of M49I over recent months in Japan compared to M49L (Supplemental Figure 2b) may be due to less favorable circulating genetic backgrounds, as most M49I sequences are associated with lineages AY.4 (714/2188 sequences) and AY.44 (172/2188), and circulated during the pre-Omicron period (Supplemental Figures 2a and 2c). These results suggest the risk of emergence of resistance associated with M49I is currently lower than for M49L, but other mutations, in the 45-49 and other regions of M^pro^, require continuous monitoring in countries where ensitrelvir is, or will soon be, approved for use.

These findings argues in favor of genetic monitoring of circulating strains to ensure that the treatments administered retain their full effectiveness. In addition, the range of antiviral treatments available against SARS-CoV-2 has become worryingly limited, with most commercial monoclonal drugs losing their activity against the most recent variants. Obviously, there is still a need to develop other antivirals targeting other viral proteins to broaden the therapeutic armory, and possibly to develop dual therapies as has been done for other viral diseases.

## Acknowledgments

We thank Magali Gilles (UVE; Marseille), Gregory Moureau (UVE; Marseille) and Camille Placidi (UVE; Marseille) for their valuable technical contribution. We thank Edyth Parker for her recommendations on SARS-CoV-2 phylogenetic inference. We thank Pr. Drosten for providing the SARSCoV-2 strain through the European Research infrastructure EVA GLOBAL.

This work was supported by the European Virus Archive Global (EVA GLOBAL) funded by the European Union’s Horizon 2020 research and innovation program under grant agreement No. 871029 and by DNDi under support by the Wellcome Trust Grant ref: 222489/Z/21/Z through the COVID-19 Therapeutics Accelerator. This work was supported by the ARBOGEN project, funded by the MSDAVENIR foundation. Part of the work was done on the Aix Marseille University antivirals platform “MaSC”.

## Author Contributions

Conceptualization : AN, FE, IS, EC and FT

Methodology :, JSD, AN, and FT

Formal analysis : SHB, JSD, RK, OB,GP,RA and FT

Investigation : SHB, JSD,RK, GP, RA, OBand FT

Resources : XdL, FT and AN

Writing original draft: SHB, JSD, AN and FT;

Writing review & editing : all co-authors

Visualization SHB, JSD, RK, OB, AN and FT

Funding acquisition LF, CM, EC, XdL, AN and FT.

