## Supplemental Data for "Generation and evaluation of protease inhibitor-resistant SARS-CoV-2 strains"

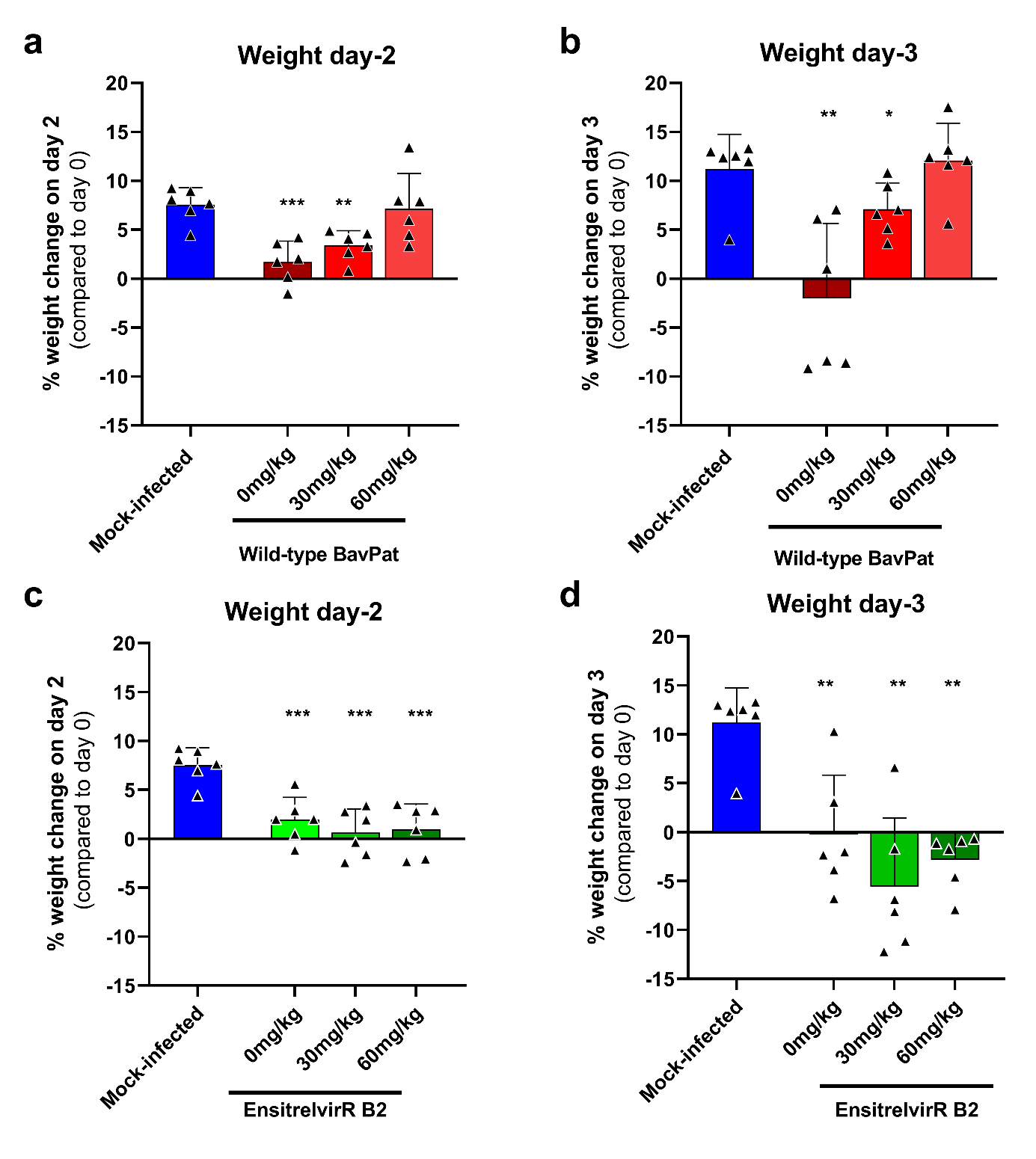

**Supplemental Figure 1**: Clinical follow up after Wild-type (Bavpat) and EnsitrelvirR B2 SARS-CoV-2 viral challenge a-b : Weight change at day 2 and at day 3 postinfection in percentage, normalized to the body weight at day 0 of Wild-type (Bavpat) infection. c-d : Weight change at day 2 and at day 3 post-infection in percentage, normalized to the body weight at day 0 of EnsitelvirR B2 infection. Bars represent means ± SD. The datas were analyzed with the unpaired t test (fig.A-C) and Mann-Whitney test (fig B-D). *P < .05, **P < .01, ***P < .001.

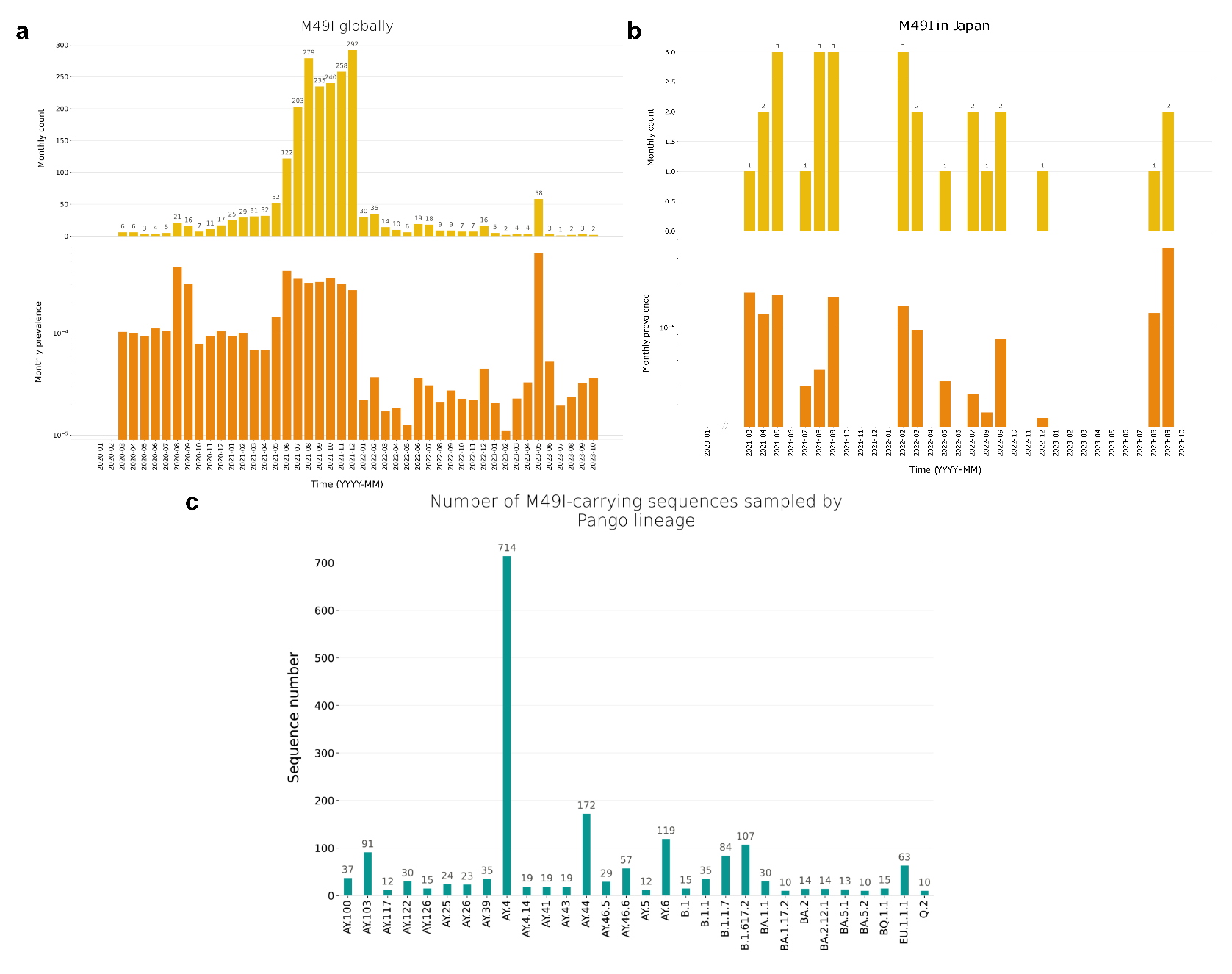
**Supplemental Figure 2**:  **M49I distribution in time, Japan, and across lineages**. a) Global monthly counts (upper panel) and prevalences (lower panel) of M49I-sequences, based on all SARS-CoV-2 genomes publicly available on GISAID as of 2023-11-13. b) Monthly counts (upper panel) and prevalences (lower panel) of M49I-sequences in Japan, based on all SARS-CoV-2 genomes publicly available on GISAID as of 2023-11-13. c) Number of M49I sequences observed by lineage.

**Supplementary Table 1:** Accession IDs, collection dates, submission dates, collection location, and Pango lineage (assigned using Pangolin lineage assigner) for all M49L sequences analyzed in this study.

**Supplementary Table 2:** GISAID accession IDs for all SARS-CoV-2 genomes (and attached metadata) in the M49L analysis.

**Data Availability**

GISAID Identifier: EPI_SET_231112rd

doi: 10.55876/gis8.231112rd

All genome sequences and associated metadata in this dataset are published in GISAID’s EpiCoV database. To

view the contributors of each individual sequence with details such as accession number, Virus name, Collection

date, Originating Lab and Submitting Lab and the list of Authors, visit 10.55876/gis8.231112rd

**Data Snapshot**

EPI_SET_231112rd is composed of 2,607 individual genome sequences.

The collection dates range from 2019-12-30 to 2023-10-30;

Data were collected in 6 countries and territories;

All sequences in this dataset are compared relative to hCoV-19/Wuhan/WIV04/2019 (WIV04), the official

reference sequence employed by GISAID (EPI_ISL_402124). Learn more at <https://gisaid.org/WIV04>.

**Supplementary Table 3**: Accession IDs, collection dates, submission dates, collection location, and Pango lineage (assigned using Pangolin lineage assigner) for all M49I sequences analyzed in this study.

**Supplementary Table** 4: GISAID accession IDs for all SARS-CoV-2 genomes (and attached metadata) in the M49I analysis.

**Data Availability**

GISAID Identifier: EPI_SET_240108hb

doi: 10.55876/gis8.240108hb

All genome sequences and associated metadata in this dataset are published in GISAID’s EpiCoV database. To

view the contributors of each individual sequence with details such as accession number, Virus name, Collection

date, Originating Lab and Submitting Lab and the list of Authors, visit 10.55876/gis8.240108hb

Data Snapshot

EPI_SET_240108hb is composed of 2,188 individual genome sequences.

The collection dates range from 2020-03-04 to 2023-10-09;

Data were collected in 55 countries and territories;

All sequences in this dataset are compared relative to hCoV-19/Wuhan/WIV04/2019 (WIV04), the official

reference sequence employed by GISAID (EPI_ISL_402124). Learn more at <https://gisaid.org/WIV04>.
